# Effects of mild winter conditions and two pyrethroid insecticides on the development and survival of juvenile common toads (*Bufo bufo*)

**DOI:** 10.1101/2023.09.06.556492

**Authors:** Andrea Kásler, Veronika Bókony, Zsanett Mikó, Dávid Herczeg, János Ujszegi, Dóra Holly, Attila Hettyey

## Abstract

Combined effects of pollutants and climate change can have long-term negative effects on Earth’s biodiversity. We tested how amphibians are affected by exposure to pesticides around metamorphosis in combination with current winter conditions or those projected to become the norm by 2100. We applied two pyrethroid insecticides, cypermethrin or deltamethrin, to newly metamorphosed common toads (*Bufo bufo*) via spraying, thereafter reared animals in outdoor enclosures, and finally exposed them either to a long&cold winter (1.5 °C for 91 days) or to a short&mild one (4.5 °C for 61 days). After hibernation, we examined changes in body mass, size and gross morphology of parotoid glands and visceral organs, and determined the phenotypic sex of the surviving toadlets. In the long&cold winter scenario, we observed high mortality rate, regardless of the chemical treatment (control: 82%, cypermethrin: 78%, deltamethrin: 70%). At the same time, mortality was significantly lower in the short&mild winter scenario (control: 33%, cypermethrin: 41%, deltamethrin: 26%). The decrease in body mass during overwintering did not differ among treatment groups. Our results contradict the hypothesis of sex-biased mortality caused by chemical pollutants, harsh or mild winter conditions or by their combination. No significant deformations of the internal organs were found in either group. Our results suggest that winter conditions becoming ever milder in the temperate zone may prove beneficial for amphibian overwinter survival, and that exposure to the tested pyrethroids at ecologically relevant concentrations after metamorphosis does not interfere with the fitness of juvenile common toads.

## 1. INTRODUCTION

The currently ongoing mass extinction of wildlife is driven by several factors, including habitat loss, overexploitation, infectious diseases, invasive species, contaminants and climate change (Blaustein et al., 2011; Collins, 2010; Shivanna, 2020; Young et al., 2016). Although these factors rarely occur in isolation and their interactive effects may often be additive or even synergistic, experimental investigations examining the combined effects of these threats are surprisingly rare.

Chemical pollution poses one of the most severe problems for both wildlife and human health (Tang et al., 2021). Pesticides are a major group of environmental pollutants, as they are used in vast quantities worldwide in pest management and vector control in agriculture and also in households (Hedlund et al., 2020; Thatheyus et al., 2013; van den Berg et al., 2012). Type II synthetic pyrethroids, like cypermethrin and deltamethrin, are among the most commonly used pesticides (van den Berg et al., 2021), because they have low levels of acute toxicity in plants, birds and mammals if compared to organochlorine and organophosphate insecticides (Katsuda, 1999). Their mode of action is via disruption of sodium channels in neurons, resulting in hypersensitivity, choreoathetosis, tremors and paralysis (Narahashi, 1996; Soderlund, 2010). Many laboratory studies have found that they are harmful to a wide range of non-target organisms, including annelids, arthropods, amphibia, fish and birds (Rehman et al., 2014), but under field conditions, their toxic effects quickly diminish due to degradation (for review see Maund et al., 2011).

Amphibian populations have gone through drastic declines worldwide (Stuart et al., 2004), and are among the most vulnerable vertebrate taxa (IUCN, 2022). In their case, exposure to pyrethroids can happen both in aquatic (as embryos or larvae) and in terrestrial environments (as juveniles and adults). Most ecotoxicological studies that examined the malign effects of pyrethroids on amphibians concentrated on aquatic life stages (Agostini et al., 2020, 2009; Berrill et al., 1993; Biga and Blaustein, 2013; David et al., 2012; Macagnan et al., 2017; Nataraj and Krishnamurthy, 2012; Vanzetto et al., 2019a). Cypermethrin and deltamethrin, two of the most heavily used pyrethroids, are often used for mosquito control, and the time and location of spraying can overlap with the emergence of juvenile amphibians from water bodies. Therefore, studies examining the effects of pyrethroids on juvenile amphibians would be important but remained scarce, although exposure to these chemicals orally or through the skin can cause histopathology and genotoxicity, or result in altered toxin profiles (Alnoaimi et al., 2021; Paetow et al., 2012; Zhou et al., 2019).

Their complex life cycle, ectothermic nature and highly permeable skin make amphibians especially vulnerable to not just chemical pollutants, but also to changes in climatic conditions. For example, changes in temperature or precipitation profiles can interfere with breeding phenology (Combes et al., 2018; While and Uller, 2014), body condition (Reading, 2007), phenotypic sex (Eggert, 2004) and disease susceptibility (Bosch et al., 2007; Cohen et al., 2017; Pounds et al., 2006; Rohr and Raffel, 2010). The fitness of amphibians may also be drastically lowered by unusual climatic conditions experienced during overwintering, which is anyway a critical time period, causing high mortality (Bradford, 1983; Clarke, 1977; Sinsch, 1988). Climate change projections predict milder and shorter winters for the temperate zone (IPCC, 2022), which may have negative effects on amphibians (Miller et al., 2018; Reading, 2007; but see Kásler et al., 2023; Üveges et al., 2016).

Here, we present an experimental test of the effects of pyrethroid pesticide exposure around metamorphosis combined with different winter conditions on the survival, body mass change and condition of hibernating juvenile common toads *Bufo bufo* (Linnaeus, 1758), a species which is widely distributed in the western Palearctic (Recuero et al., 2012). Common toads may benefit from mild winters (Üveges et al., 2016), but stressors, such as pesticide exposure may decrease their ability to cope with changing hibernation conditions. We used juvenile toads reared from the egg stage under seminatural conditions, performed cutaneous pyrethroid exposure shortly after the completion of metamorphosis, and exposed individuals to winter conditions that are currently typical for the study area (1.5°C for 91 days Sinsch, 1988; van Gelder et al., 1986) or to conditions that are projected to become the norm by around 2100 (4.5°C for 61 days; IPCC, 2022).

## 2. MATERIALS AND METHODS

### 2.1 Chemicals

We obtained cypermethrin and deltamethrin from Sigma-Aldrich (CAS numbers 52315-07-8 and 52918-63-5, respectively). Stock solutions were prepared by dissolving 50 mg cypermethrin in 2 ml 96 % ethanol, and 60 mg deltamethrin in 4 mL 96 % ethanol. For the exposure we diluted 3 microliters of the cypermethrin stock and 13.5 microliters of the deltamethrin stock in 300 mL RSW (reconstituted soft water; USEPA, 2002) We chose the final concentrations of both active ingredients to accord to permits issued for two commercial insecticide formulations [Cyperkill Max (Arysta LifeScience Magyarország Ltd., Hungary): 0.75 mg cypermethrin / m^2^; Aqua K-Othrine (Bayer Hungaria Ltd., Hungary): 0.1 mg deltamethrin / m^2^]. For the control group we diluted 540 microliters of 96 % EtOH in 300 ml RSW.

### 2.2 Collection and rearing of animals

On 7 April 2021 we collected 200 eggs from each of 18 egg-strings of the common toad. Eggs originated from three ponds in the Pilis-Visegrádi Mountains, Hungary (from six egg-strings in the Békás-tó: 47°34ʹ34.72ʺN, 18°52ʹ8.06ʺE, four egg-strings in the Ilona-tó: 47°42ʹ47.7ʺN, 19°02ʹ25.8ʺE, and eight egg-strings in the Jávor-tó: 47°42ʹ49.7ʺN, 19°01ʹ11.1ʺE). We transported the eggs to the Experimental Station Júliannamajor of the Plant Protection Institute, Centre for Agricultural Research, Budapest. We kept eggs separated by egg-string (family hereafter) in plastic containers (32×22×16 cm) filled with 0.7 L reconstituted soft water (RSW) at a constant temperature of 16 °C and a light:dark cycle adjusted weekly to the conditions outside. Nine days after hatching, when all larvae reached development stage 25 (Gosner, 1960), we placed 50 tadpoles separated by family into outdoor mesocosms. We set up mesocosms in March by filling plastic tubs (85×57×51 cm) with 130 liters of aged tap water and adding 50 g dried beech (*Fagus sylvatica*) leaves to provide tadpoles with nutrients and refuges. To enhance algal growth and start up a self-sustaining ecosystem, we inoculated mesocosms with 1 liter of pond water containing bacterio-, phyto- and zooplankton. Four weeks later we added another 600 ml pond water to each mesocosm to boost zooplankton density and thereby reduce algal bloom. We covered mesocosms with mosquito screen lids to prevent colonization by predatory insects. When the first individuals reached development stage 42 (emergence of forelimbs, start of metamorphosis), we removed leaves and monitored mesocosms daily for metamorphosing individuals, captured them using small dip-nets and placed them into transparent plastic boxes (52×35×25 cm; one box for each mesocosm) covered with a perforated lid and placed into a shady outdoor area. Boxes contained approximately 1.5 litres of mesocosm water and were slightly tilted to provide both a water-covered as well as a dry area. When individuals completed metamorphosis (development stage 46), we chose 14 families exhibiting high survival rates and placed animals separated by family into plastic containers (60×40×30 cm; 50 individuals per container) lined with wet paper towels as a substrate, and containing a large piece of egg carton to provide shelters. Twice a week we sprinkled boxes with RSW to maintain humidity and fed metamorphs with commercially acquired small crickets (*Acheta domesticus*, instar stage 1–2).

On 14 July, we assigned nine groups of toadlets into boxes by randomly selecting four individuals from each family, thus each box contained 56 individuals. We previously lined the boxes with sterilized soil. We sprayed the toadlets with the two pesticides and the ethanol control using hand sprayers, with the entire content of sprayers emptied into each box. Eight hours later, we removed animals from the boxes and placed them into outdoor enclosures located in a forested area at the experimental station. We kept animals separated by treatment. Enclosures measured 3 × 3 m and were delimited by fences constructed of fine-meshed mosquito net. Fences were 60 cm high and were dug 20 cm into the soil. Enclosures were covered with chicken wire to allow small invertebrates to enter from above and serve as food for toadlets, but keep out predators. We equipped each enclosure with two plastic poultry waterers to provide water permanently. We checked enclosures weekly for incidental damage and refilled waterers if necessary. Every 2 weeks we supplemented naturally occurring food by releasing into each enclosure ca. 500 small crickets (*A. domesticus*; instar stage 2–3). On 13 and 15 October, we captured surviving individuals by carefully searching enclosures twice (Table S1).

### 2.3 Overwintering and dissection of juveniles

Due to limited space we used a total of 162 toadlets for overwintering, 54 individuals from each chemical treatment (TABLE S1). On 18 October we measured the body mass of individuals (± 0.1 mg) and placed them individually into 50 ml centrifuge tubes filled with 20 ml of a 1:1 mixture of sterilized and dried soil and sand, moistened with 5 ml aged tap water to provide a substrate and prevent desiccation (for further details see Üveges et al., 2016). We covered one end of tubes with mosquito nets to allow air exchange and easy watering but at the same time prevent animals from escaping. We assigned juveniles from all three treatment groups randomly to one of two hibernation scenarios while keeping body masses balanced among the six treatment groups (LM; F = 0.006, df = 2, *P* = 0.99; N = 27 / group). We placed tubes into two laboratory refrigerators with forced air convection (Pol-Eko-Aparatura, Wodzisław Śląski, Poland) set to 10 °C. Two days later, we lowered the temperature to 7.5 °C and maintained it for three weeks to simulate transient autumn temperatures (FIGURE 1). We initiated hibernation on 10 November by lowering the temperature to 4.5 °C in the short&mild (SM) and to 1.5 °C in the long&cold (LC) winter scenario, respectively. We kept track of actual temperatures every five minutes using the refrigerators’ inbuilt thermometers. Mean temperature ± SE during the SM and the LC treatments were 4.495 ± 0.004 and 1.486 ± 0.003 °C, respectively. We sprinkled aged tap water into tubes twice a week to prevent desiccation. At these occasions we checked if all individuals were alive, and recorded mortalities. Sixty-one (SM treatment) and 91 days (LC treatment) after the start of hibernation we raised the temperature to 7.5 °C in the refrigerators. Animals were kept at this temperature for seven days. Finally, we measured body mass of individuals and humanely euthanized them using a water bath containing 6.6 g/L MS-222 buffered to neutral pH with the same amount of Na2HPO4.

**Figure 1.**
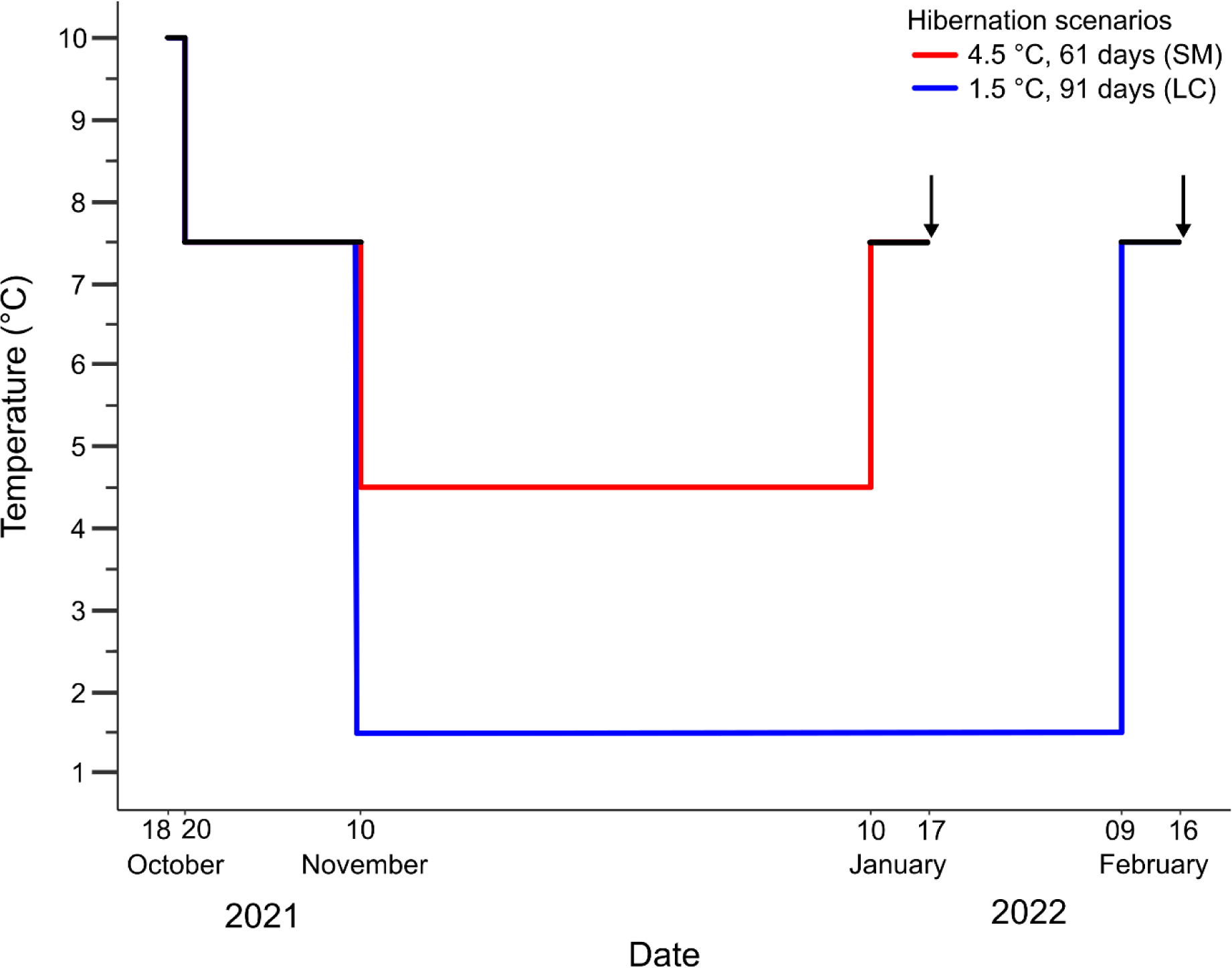
A schematic representation of the applied overwintering scenarios. Arrows show the dates of weighing after hibernation. SM = short&mild winter; LC = long&cold winter.

After dissection, we examined the parotoid glands and the internal organs under an Apomic SHD 200 digital microscope (Apokromat Ltd., Hungary) and recorded abnormalities. We measured the length and width of parotoid glands using the microscope’s measuring tool and estimated their area assuming the shape of an ellipse (Regueira et al., 2017). For analysis, we averaged the areas of the two parotoids of each toadlet. We recorded the quantity of fat bodies (no/few, normal or extreme), and whether the individual had testes (male) or ovaries (female), or abnormally looking gonads (uncertain sex). We photographed the spleen of the individuals and measured the area of the spleen and the percentage area of its pigmented spots with ImageJ. The measurements were converted from pixels to mm^2^ by photographing a size standard at the same magnification as the spleens.

### 2.4 Statistical analyses

All statistical analyses were conducted in R (version 4.2.2, “R Core Team,” 2020). In all models, we entered pre-hibernation body mass, winter scenario, pesticide treatment, and the latter two’s interaction as independent variables. We analyzed the survival of toadlets during hibernation using Cox’s proportional hazards model (‘coxph’). To analyze the differences in the overall proportion of toadlets that survived until dissection between treatment groups, we used a generalized linear model (‘glm’ with binomial link function). We did not analyze differences of survival between the outdoor enclosures, as sample sizes were very low (two or three enclosures per treatment). To test for sex-dependent mortality we compared the number of phenotypic males and females of the two pesticide treatment groups to the control group of both winter scenarios using Fisher’s exact test. We did not apply a more sophisticated analysis, because the overall sample size of toadlets surviving to phenotypic sexing was low, with zero individuals in certain combinations of treatment by sex. In case of continuous dependent variables (change in body mass, parotoid size, spleen size, and spleen pigmentation), we fitted linear models. In case of the quantity of fat reserves, we fitted a proportional odds model (‘clm’ with logit link function), entering the winter scenario, pesticide treatment and their interaction as independent variables.

## 3. RESULTS

### 3.1 Survival

Spraying the individuals with the pyrethroids and the ethanol control did not cause acute mortality. Overall, 73 toadlets out of 162 survived until dissection (N = 54 in the SM group, and N = 19 in the LC group; TABLE 1). The winter scenario significantly affected the survival of animals: individuals in the short&mild winter group had three times higher survival probability than individuals in the long&cold winter group (COXPH; χ^2^1 = 11.39, *P* < 0.001; FIGURE 2, TABLE 1). At the same time, pre-hibernation body mass also had a significant effect on survival probability: juveniles with greater pre-hibernation body mass had a lower hazard of dying (χ^2^1 = 15.43, *P* < 0.001; mean pre-hibernation body mass ± S.E. for survivors: 564.84 ± 12.64 mg, for those that died: 488.03 ± 10.54 mg). The effects of pesticide treatments (χ^2^2 = 0.71, *P* = 0.70) and the interaction of winter scenario and pesticide treatment (χ^2^2 = 1.94, *P* = 0.38) were non-significant.

**TABLE 1:** The number of toadlets that died during hibernation (hib.) or between hibernation and dissection (dis.), and the number of toadlets that survived until dissection in the long&cold (LC) and in the short&mild (SM) winter scenarios.

| Treatment | LC winter |  |  | SM winter |  |  |
| --- | --- | --- | --- | --- | --- | --- |
|  | Died during hib. | Died between hib. and dis. | Survived until dis. | Died during hib. | Died between hib. and dis. | Survived until dis. |
| Control | 22 | 0 | 5 | 3 | 6 | 18 |
| Cypermethrin | 20 | 1 | 6 | 5 | 6 | 16 |
| Deltamethrin | 17 | 2 | 8 | 5 | 2 | 20 |

**Figure 2.**
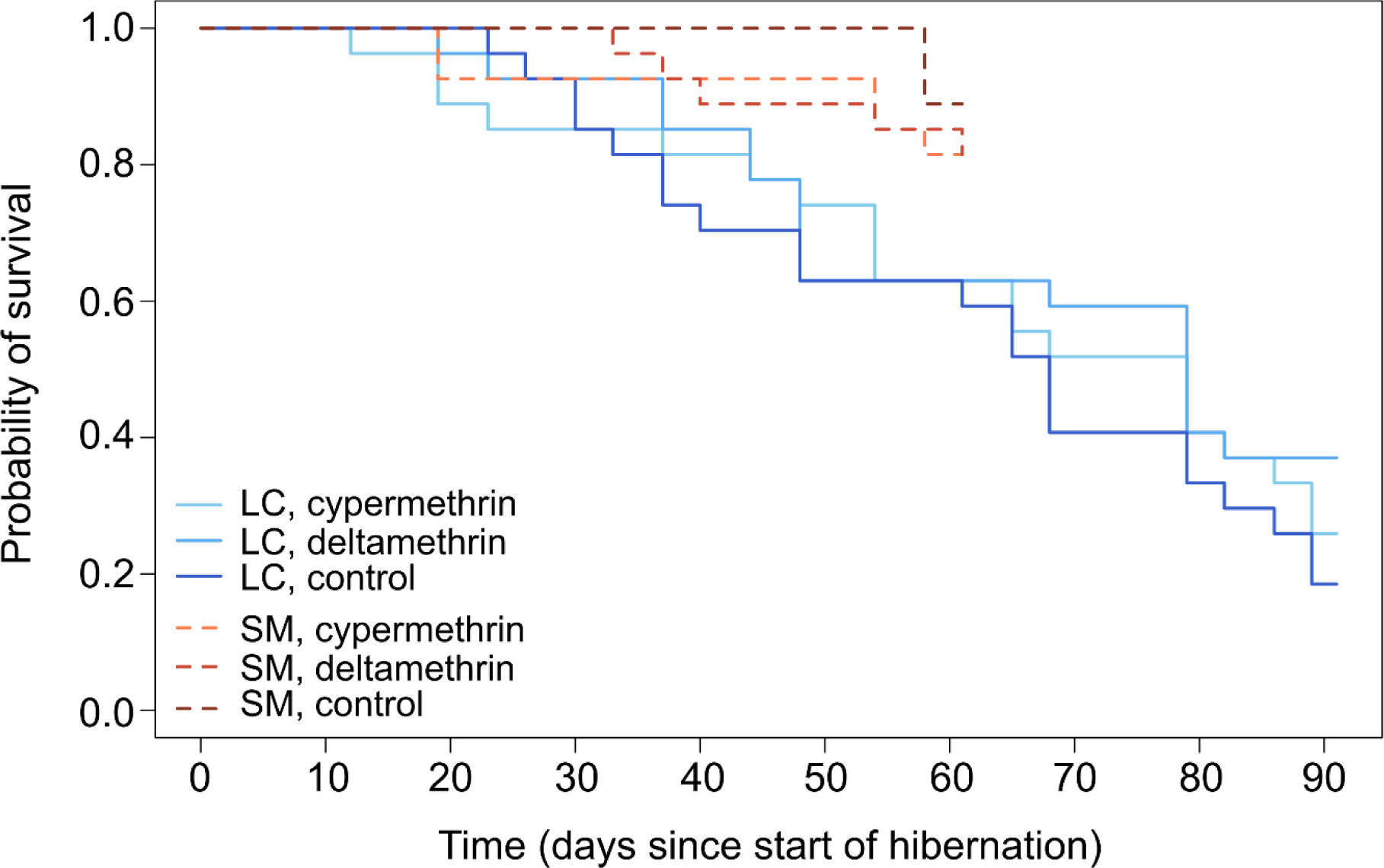
Survival probabilities of juvenile common toads during hibernation in different treatment combinations visualized by Kaplan-Meier curves. LC = long&cold winter scenario; SM = short&mild winter scenario.

A total of 17 toadlets died during the week between the end of hibernation and dissection: three in the SM and fourteen in the LC groups (TABLE 1). The number of toadlets that died before dissection showed a similar pattern to those that died during hibernation: significantly more toadlets died in the LC groups than in the SM groups (GLM; χ^2^1 = 36.56, *P* < 0.001), but pesticide treatment alone (χ^2^2 = 0.84, *P* = 0.658) or in interaction with the winter scenarios (χ^2^2 = 0.37, *P* = 0.833) did not have a significant effect.

The sex ratio of surviving toadlets did not differ between the treatment groups in neither winter scenario (Fisher’s exact test, *P* = 0.933 and *P* = 0.266 for LC and SM groups, respectively; TABLE 2).

**TABLE 2:** The phenotypic sex of dissected individuals in treatment combinations. We found three individuals with uncertain sex based on the macroscopic characteristics of gonads, but because of the low incidence, we did not follow up on this.

| Treatment | LC winter |  |  | SM winter |  |  |
| --- | --- | --- | --- | --- | --- | --- |
|  | Female | Male | Uncertain | Female | Male | Uncertain |
| Control | 2 | 3 | 0 | 6 | 12 | 1 |
| Cypermethrin | 4 | 2 | 0 | 10 | 7 | 1 |
| Deltamethrin | 4 | 3 | 1 | 10 | 10 | 0 |

### 3.2 Body mass loss

Winter scenario (F = 0.26, df = 1, *P* = 0.613), pesticide treatment (F = 2.52, df = 2, *P* = 0.089), or their interaction (F = 0.87, df = 2, *P* = 0.423) did not have a significant effect on body mass loss during overwintering (TABLE 3).

**TABLE 3:** mean ± SE of each outcome variable in each treatment group estimated from the models. Contr. = control; Cyp. = cypermethrin; Delt. = deltamethrin.

| Outcome variable | LC winter |  |  | SM winter |  |  |
| --- | --- | --- | --- | --- | --- | --- |
|  | Contr. | Cyp. | Delt. | Contr. | Cyp. | Delt. |
| Change in body mass (mg) | -58.2 ± 14.7 | -38.6 ± 13.5 | -52.7 ± 12.1 | -35.8 ± 7.74 | -37.2 ± 8.25 | -58.5 ± 7.37 |
| Mean area of parotoid glands (mm<sup>2</sup>) | 3.92 ± 0.36 | 3.74 ± 0.32 | 4.23 ± 0.28 | 3.90 ± 0.19 | 3.93 ± 0.19 | 3.71 ± 0.18 |
| Spleen size (mm<sup>2</sup>) | 0.44 ± 0.05 | 0.31 ± 0.05 | 0.30 ± 0.04 | 0.41 ± 0.03 | 0.42 ± 0.03 | 0.38 ± 0.03 |
| Spleen pigmentation (%) | 1.91 ± 1.26 | 2.17 ± 1.13 | 1.51 ± 0.99 | 2.97 ± 0.69 | 2.65 ± 0.65 | 3.21 ± 0.58 |

### 3.3 Morphology

The size of the parotoid glands, the size of the spleen and the proportion of pigmented areas in the spleen were not affected by neither coefficients, except that pre-hibernation body mass had a positive effect on the size of the parotoid glands and the spleen (TABLE 3 and 4). The quantity of fat reserves was also not affected by the examined variables (FIGURE S1).

**Table 4:**
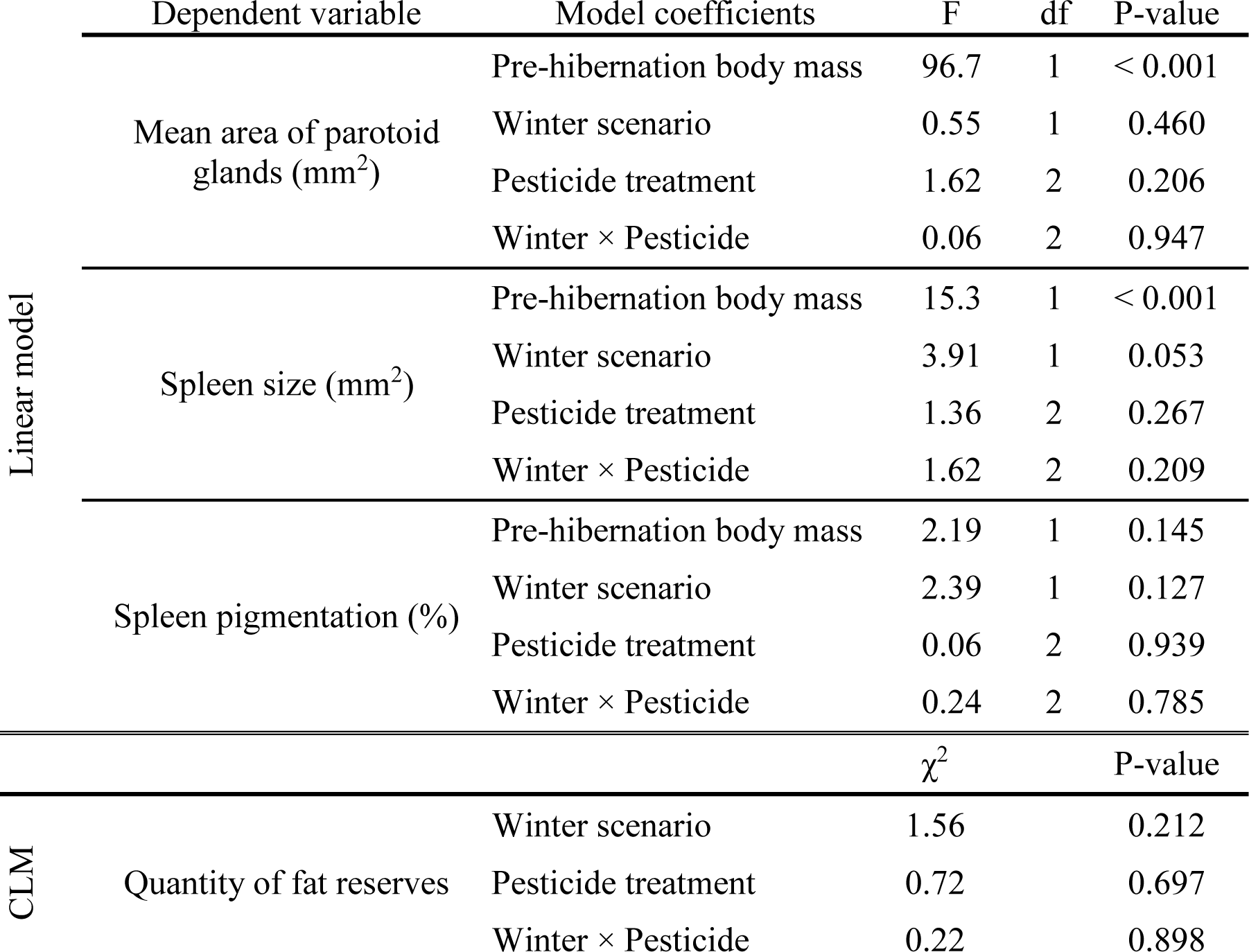
Coefficients of LM and CLM models for size of the parotoid glands and spleen, spleen pigmentation and quantity of fat reserves. CLM = cumulative link model.

## 4. DISCUSSION

Our experiment revealed no harmful effects of pyrethroid exposure around metamorphosis on overwintering juvenile common toads either at currently typical winter conditions or at milder ones projected for 2100. We found no effect of the tested pesticides, cypermethrin and deltamethrin on the survival of individuals during hibernation. We speculate that this could be the result of the appliance of only a short, acute exposure around metamorphosis to ecologically relevant concentrations of the pesticides. We know of no other study that examined the long-term effects of cypermethrin- or deltamethrin exposure applied after metamorphosis on the survival of amphibians. However, experiments reporting toxicity of various pyrethroids on aquatic life stages of amphibians mostly applied chronic exposure (several days) and/or very high concentrations of the chemicals (Agostini et al., 2009; Berrill et al., 1993; Biga and Blaustein, 2013; Macagnan et al., 2017; Vanzetto et al., 2019b). Experiments regarding the survival of metamorphic amphibians exposed to pesticides are scarce, also rely on chronic exposures (Jones et al., 2017; Zhou et al., 2019). It is important to note that our study does not allow conclusions on potential effects of repeated or longer exposure to the pesticides, or on how spraying at times other than shortly after metamorphosis affects juvenile survival.

We found that survival probability was higher in the short&mild than in the long&cold winter group. This finding is similar to that described in Üveges et al. (2016), while Garner et al. (2011) found no relationship between the survival of juvenile common toads and different overwintering regimes. Our results suggest, that shorter and milder winters, which are predicted to increasingly become the norm over the course of the 21^st^ century, may benefit common toad populations in the temperate zone, but we have to point out that common toads have a wide distribution in Eurasia, reaching from the Mediterranean beyond the northern Arctic Circle, and populations with different levels of cold-adaptation might show diverse sensitivity to milder winter conditions. On the other hand, we observed higher mortality rates (60.5% overall) than was reported in previous experiments (12.4% and 1.9% overall, respectively). This may have occurred partly because the pre-hibernation body mass of toadlets was greater in both experiments (approximately two times greater in Üveges et al., and one and a half times greater in Garner et al.), suggesting a better body condition of individuals, and greater pre-hibernation body mass indicates better survival of hibernating amphibians (Altwegg and Reyer, 2003; Üveges et al., 2016). On the other hand, Garner et al. used higher temperatures during hibernation (8 °C and 4 °C for different period-lengths in the normal and changing climate groups), which may also have contributed to the lower mortality rate. Another laboratory experiment with juvenile *Rana dalmatina* also found no difference in survival between different hibernation regimes (Kásler et al., 2023), while a field study found significant relationship between the occurrence of mild winters and decline in the body condition of female common toads.

Our results showed the absence of abnormal gonads and skewed sex ratios of surviving toadlets. Gonadal sex differentiation of common toads starts relatively late compared to other amphibians (Falconi et al., 2004; Ogielska and Kotusz, 2004), so we expected to find several individuals with uncertain sex and abnormal gonads as we applied the pesticides shortly after metamorphosis, i.e. during the sensitive period. Nonetheless, the low number of individuals with uncertain sex and the lack of highly skewed sex ratios in the pesticide-exposed groups indicate that acute exposure to cypermethrin and deltamethrin in low concentrations around metamorphosis do not cause severe alterations in sexual differentiation of the gonads nor do they cause striking sex-specific mortality.

Our result that changes in body mass were similar across animals exposed to different overwintering conditions is in accordance with previous studies (Garner et al., 2011; Kásler et al., 2023). Nonetheless, Üveges et al. (2016) found that longer and colder winter regimes can cause greater decreases in body mass during hibernation. Future studies should conduct more direct experiments regarding the effects of climate change and pesticide exposure on the body condition of amphibians.

We found no difference in the size of the parotoid glands between pesticide treatment groups and winter scenarios, suggesting that pyrethroid exposure and milder winters did not heavily influence the toxin production of individuals. Parotoid glands contain most of the toad’s bufadienolides, and larger parotoid glands produce more toxins (Bókony et al., 2019; Llewelyn et al., 2012). Thus, a considerable decrease in toxin production can result in smaller parotoid glands. Further analysis of the composition of the venom would be important in future studies, as Zhou et al. (2019) found that a high level of cutaneous deltamethrin contamination can lead to a marked decrease in the bufadienolide content in toad venom, which may result in lowered protection against predators.

We found no major long-term effects of pesticide exposure and milder winter conditions on fat reserves and spleen morphology. Fat bodies are essential for winter survival and reproduction after hibernation, thus further experiments should investigate whether pesticides and altered winter conditions have an effect on their quantity. Although size and pigmentation of the spleen are good indicators of immune function and pathogen resistance in ectothermic vertebrates (Hadidi et al., 2008; Steinel and Bolnick, 2017), we found no information on the morphological changes of the spleen overwinter. Our result regarding the lack of differences between treatment groups does not support the hypothesis that pesticide exposure or altered winter conditions severely impaired the immune function of animals.

In summary, our results showed that cutaneous exposure to cypermethrin or deltamethrin in ecologically relevant quantities, even if followed by harsh or unusually mild winter conditions, does not harm juvenile post-metamorphic common toads. These results draw attention to the fact that strict regulations of pesticide use are essential and may indeed be effective for avoiding malign effects to wildlife. It is clear that amphibian species can differ in their sensitivity to pollutants, climate change and many other factors causing stress, while it is also safe to assume that they vary in their vulnerability to various combinations of these factors (Blaustein and Kiesecker, 2002), leaving a vast space for future studies on the joint effects of multiple stressors on amphibians.

## AUTHOR CONTRIBUTIONS

A.K. and A.H. conceived and designed the experiment. A.K., V.B., Z.M., D.He., J.U. and D.Ho. performed the experiment. A.K. analyzed the data and wrote the first draft of the manuscript, while all other authors contributed to the final version.

## ACKNOWLEDGEMENTS

We thank M. Szederkényi for assistance during the experiment, and N. Ujhegyi for help in capturing the juvenile toadlets. All experimental procedures were approved by the Ethical Commission of the Plant Protection Institute and carried out according to the permits issued by the Government Agency of Pest County (Department of Environmental Protection and Nature Conservation, PE-06/KTF/8060-1/2018, PE-06/KTF/8060-2/2018, PE-06/KTF/8060-3/2018 and PE/EA/295-7/2018).

## REFERENCES

1. Agostini, M.G., Natale, G.S., Ronco, A.E., 2009. Impact of endosulphan and cypermethrin mixture on amphibians under field use for biotech soya bean production. International Journal of Environment and Health 3, 379–389. 10.1504/IJENVH.2009.030109

2. Agostini, M.G., Roesler, I., Bonetto, C., Ronco, A.E., Bilenca, D., 2020. Pesticides in the real world: The consequences of GMO-based intensive agriculture on native amphibians. Biological Conservation 241, 108355. 10.1016/j.biocon.2019.108355

3. Alnoaimi, F., Dane, H., Şişman, T., 2021. Histopathologic and genotoxic effects of deltamethrin on marsh frog, *Pelophylax ridibundus* (Anura: Ranidae). Environmental Science and Pollution Research 28, 3331–3343. 10.1007/s11356-020-10711-5

4. Altwegg, R., Reyer, H.U., 2003. Patterns of natural selection on size at metamorphosis in water frogs. Evolution 57, 872–882. 10.1111/j.0014-3820.2003.tb00298.x

5. Berrill, M., Bertram, S., Wilson, A., Louis, S., Brigham, D., Stromberg, C., 1993. Lethal and Sublethal Impacts of Pyrethroid Insecticides on Amphibian Embryos and Tadpoles. Environmental Toxicology and Chemistry 12, 525. 10.1897/1552-8618(1993)12[525:lasiop]2.0.co;2

6. Biga, L.M., Blaustein, A.R., 2013. Variations in lethal and sublethal effects of cypermethrin among aquatic stages and species of anuran amphibians. Environmental Toxicology and Chemistry 32, 2855–2860. 10.1002/etc.2379

7. Blaustein, A.R., Han, B.A., Relyea, R.A., Johnson, P.T.J., Buck, J.C., Gervasi, S.S., Kats, L.B., 2011. The complexity of amphibian population declines: Understanding the role of cofactors in driving amphibian losses. Annals of the New York Academy of Sciences 1223, 108–119. 10.1111/j.1749-6632.2010.05909.x

8. Blaustein, A.R., Kiesecker, J.M., 2002. Complexity in conservation : Lessons from the global decline of amphibian populations. Ecology letters 5, 597–608.

9. Bókony, V., Üveges, B., Verebélyi, V., Ujhegyi, N., Móricz, Á.M., 2019. Toads phenotypically adjust their chemical defences to anthropogenic habitat change. Scientific Reports 9, 1–8. 10.1038/s41598-019-39587-3

10. Bosch, J., Carrascal, L.M., Durán, L., Walker, S., Fisher, M.C., 2007. Climate change and outbreaks of amphibian chytridiomycosis in a montane area of Central Spain; is there a link? Proceedings of the Royal Society B: Biological Sciences 274, 253–260. 10.1098/rspb.2006.3713

11. Bradford, D.F., 1983. Winterkill, oxygen relations, and energy metabolism of a submerged dormant amphibian, Rana muscosa. Ecology 64, 1171–1183.

12. Clarke, R.D.., 1977. Postmetamorphic Survivorship of Fowler’s Toad, *Bufo woodhousei fowleri*. Copeia 3, 594–597.

13. Cohen, J.M., Venesky, M.D., Sauer, E.L., Civitello, D.J., McMahon, T.A., Roznik, E.A., Rohr, J.R., 2017. The thermal mismatch hypothesis explains host susceptibility to an emerging infectious disease. Ecology Letters 20, 184–193. 10.1111/ele.12720

14. Collins, J.P., 2010. Amphibian decline and extinction: What we know and what we need to learn. Diseases of Aquatic Organisms 92, 93–99. 10.3354/dao02307

15. Combes, M., Pinaud, D., Barbraud, C., Trotignon, J., Brischoux, F., 2018. Climatic influences on the breeding biology of the agile frog (Rana dalmatina). Science of Nature 105. 10.1007/s00114-017-1530-0

16. David, M., Marigoudar, S.R., Patil, V.K., Halappa, R., 2012. Behavioral, morphological deformities and biomarkers of oxidative damage as indicators of sublethal cypermethrin intoxication on the tadpoles of D. melanostictus (Schneider, 1799). Pesticide Biochemistry and Physiology 103, 127–134. 10.1016/j.pestbp.2012.04.009

17. Eggert, C., 2004. Sex determination : the amphibian models 44, 539–549. 10.1051/rnd:2004062

18. Falconi, R., Dalpiaz, D., Zaccanti, F., 2004. Ultrastructural aspects of gonadal morphogenesis in Bufo bufo (Amphibia Anura) 1. Sex differentiation. Journal of Experimental Zoology Part A: Comparative Experimental Biology 301, 378–388. 10.1002/jez.a.20069

19. Garner, T.W.J., Rowcliffe, J.M., Fisher, M.C., 2011. Climate change, chytridiomycosis or condition: An experimental test of amphibian survival. Global Change Biology 17, 667– 675. 10.1111/j.1365-2486.2010.02272.x

20. Gosner, K.L., 1960. A Simplified Table for Staging Anuran Embryos Larvae with Notes on Identification. Herpetoligica 16, 183–190. 10.2307/3890061

21. Hadidi, S., Glenney, G.W., Welch, T.J., Silverstein, J.T., Wiens, G.D., 2008. Spleen Size Predicts Resistance of Rainbow Trout to Flavobacterium psychrophilum Challenge . The Journal of Immunology 180, 4156–4165. 10.4049/jimmunol.180.6.4156

22. Hedlund, J., Longo, S.B., York, R., 2020. Agriculture, Pesticide Use, and Economic Development: A Global Examination (1990–2014). Rural Sociology 85, 519–544. 10.1111/ruso.12303

23. IPCC, 2022. Chapter 13: Europe, in: Climate Change 2022: Impacts, Adaptations and Vulnerability. p. 143.

24. IUCN, 2022. The IUCN red list of threatened species. Version 2022–2.

25. Jones, D.K., Dang, T.D., Urbina, J., Bendis, R.J., Buck, J.C., Cothran, R.D., Blaustein, A.R., Relyea, R.A., 2017. Effect of Simultaneous Amphibian Exposure to Pesticides and an Emerging Fungal Pathogen, Batrachochytrium dendrobatidis. Environmental Science and Technology 51, 671–679. 10.1021/acs.est.6b06055

26. Kásler, A., Holly, D., Herczeg, D., Ujszegi, J., Hettyey, A., 2023. Chytridiomycosis and climate change: exposure to *Batrachochytrium dendrobatidis* and mild winter conditions do not increase mortality in juvenile agile frogs during hibernation. Animal Conservation 1–9. 10.1111/acv.12851

27. Katsuda, Y., 1999. Development of and future prospects for pyrethroid chemistry. Pesticide Science 55, 775–782.

28. Llewelyn, J., Bell, K., Schwarzkopf, L., Alford, R.A., Shine, R., 2012. Ontogenetic shifts in a prey’s chemical defences influence feeding responses of a snake predator. Oecologia 169, 965–973. 10.1007/s00442-012-2268-1

29. Macagnan, N., Rutkoski, C.F., Kolcenti, C., Vanzetto, G. V., Macagnan, L.P., Sturza, P.F., Hartmann, P.A., Hartmann, M.T., 2017. Toxicity of cypermethrin and deltamethrin insecticides on embryos and larvae of Physalaemus gracilis (Anura: Leptodactylidae). Environmental Science and Pollution Research 24, 20699–20704. 10.1007/s11356-017-9727-5

30. Maund, S.J., Campbell, P.J., Giddings, J.M., Hamer, M.J., Henry, K., Pilling, E.D., Warinton, J.S., Wheeler, J.R., 2011. Ecotoxicology of Synthetic Pyrethroids, in: Matsuo, N., Mori, T. (Eds.), Pyrethroids. Topics in Current Chemistry, Vol 314. Springer Berlin Heidelberg, pp. 137–165. 10.1007/128_2011_260

31. Miller, D.A.W., Grant, E.H.C., Muths, E., Amburgey, S.M., Adams, M.J., Joseph, M.B., Waddle, J.H., Johnson, P.T.J., Ryan, M.E., Schmidt, B.R., Calhoun, D.L., Davis, C.L., Fisher, R.N., Green, D.M., Hossack, B.R., Rittenhouse, T.A.G., Walls, S.C., Bailey, L.L., Cruickshank, S.S., Fellers, G.M., Gorman, T.A., Haas, C.A., Hughson, W., Pilliod, D.S., Price, S.J., Ray, A.M., Sadinski, W., Saenz, D., Barichivich, W.J., Brand, A., Brehme, C.S., Dagit, R., Delaney, K.S., Glorioso, B.M., Kats, L.B., Kleeman, P.M., Pearl, C.A., Rochester, C.J., Riley, S.P.D., Roth, M., Sigafus, B.H., 2018. Quantifying climate sensitivity and climate-driven change in North American amphibian communities. Nature Communications 9, 1–15. 10.1038/s41467-018-06157-6

32. Narahashi, T., 1996. Neuronal ion channels as the target site of insecticides. Pharmacology and Toxicology 79, 1–14. 10.1111/j.1600-0773.1996.tb00234.x

33. Nataraj, M.B., Krishnamurthy, S. V., 2012. Effects of combinations of malathion and cypermethrin on survivability and time of metamorphosis of tadpoles of Indian cricket frog (Fejervarya limnocharis). Journal of Environmental Science and Health - Part B Pesticides, Food Contaminants, and Agricultural Wastes 47, 67–73. 10.1080/03601234.2012.611428

34. Ogielska, M., Kotusz, A., 2004. Pattern and Rate of Ovary Differentiation with Reference to Somatic Development in Anuran Amphibians. Journal of Morphology 259, 41–54. 10.1002/jmor.10162

35. Paetow, L.J., Daniel McLaughlin, J., Cue, R.I., Pauli, B.D., Marcogliese, D.J., 2012. Effects of herbicides and the chytrid fungus Batrachochytrium dendrobatidis on the health of post-metamorphic northern leopard frogs (Lithobates pipiens). Ecotoxicology and Environmental Safety 80, 372–380. 10.1016/j.ecoenv.2012.04.006

36. Pounds, J.A., Bustamante, M.R., Coloma, L.A., Consuegra, J.A., Fogden, M.P.L., Foster, P.N., La Marca, E., Masters, K.L., Merino-Viteri, A., Puschendorf, R., Ron, S.R., Sánchez-Azofeifa, G.A., Still, C.J., Young, B.E., 2006. Widespread amphibian extinctions from epidemic disease driven by global warming. Nature 439, 161–167. 10.1038/nature04246

37. R Core Team [WWW Document], 2020. . R: A language and environment for statistical computing. R Foundation for Statistical Computing, Vienna, Austria. URL https://www.r-project.org/

38. Reading, C.J., 2007. Linking global warming to amphibian declines through its effects on female body condition and survivorship. Oecologia 151, 125–131. 10.1007/s00442-006-0558-1

39. Recuero, E., Canestrelli, D., Vörös, J., Szabó, K., Poyarkov, N.A., Arntzen, J.W., Crnobrnja- Isailovic, J., Kidov, A.A., Cogâlniceanu, D., Caputo, F.P., Nascetti, G., Martínez-Solano, I., 2012. Multilocus species tree analyses resolve the radiation of the widespread Bufo bufo species group (Anura, Bufonidae). Molecular Phylogenetics and Evolution 62, 71–86. 10.1016/j.ympev.2011.09.008

40. Regueira, E., Dávila, C., Sassone, A.G., O’Donohoe, M.E.A., Hermida, G.N., 2017. Post- metamorphic development of skin glands in a true toad: Parotoids versus dorsal skin. Journal of Morphology 278, 652–664. 10.1002/jmor.20661

41. Rehman, H., Aziz, A.T., Saggu, S., Abbas, Z.K., Mohan, A., Ansari, A.A., 2014. Systematic review on pyrethroid toxicity with special reference to deltamethrin. Journal of Entomology and Zoology Studies 2, 60–70.

42. Rohr, J.R., Raffel, T.R., 2010. Linking global climate and temperature variability to widespread amphibian declines putatively caused by disease. Proceedings of the National Academy of Sciences of the United States of America 107, 8269–8274. 10.1073/pnas.0912883107

43. Shivanna, K.R., 2020. The Sixth Mass Extinction Crisis and its Impact on Biodiversity and Human Welfare. Resonance 25, 93–109. 10.1007/s12045-019-0924-z

44. Sinsch, U., 1988. Seasonal changes in the migratory behaviour of the toad *Bufo bufo*: direction and magnitude of movements. Oecologia 76, 390–398. 10.1007/BF00377034

45. Soderlund, D.M., 2010. Toxicology and Mode of Action of Pyrethroid Insecticides, in: Krieger, R. (Ed.), Hayes’ Handbook of Pesticide Toxicology. Academic Press, London, pp. 1665–1686.

46. Steinel, N.C., Bolnick, D.I., 2017. Melanomacrophage centers as a histological indicator of immune function in fish and other poikilotherms. Frontiers in Immunology 8, 1–8. 10.3389/fimmu.2017.00827

47. Stuart, S.N., Chanson, J.S., Cox, N.A., Young, B.E., Rodrigues, A.S.L., Fischman, D.L., Waller, R.W., 2004. Status and trends of amphibian declines and extinctions worldwide. Science 306, 1783–1786. 10.1126/science.1103538

48. Tang, F.H.M., Lenzen, M., McBratney, A., Maggi, F., 2021. Risk of pesticide pollution at the global scale. Nature Geoscience 14, 206–210. 10.1038/s41561-021-00712-5

49. Thatheyus, A.J., Deborah, A., Selvam, G., 2013. Synthetic Pyrethroids: Toxicity and Biodegradation. Applied Ecology and Environmental Sciences 1, 33–36. 10.12691/aees-1-3-2

50. USEPA, 2002. Dilution water, in: Methods for Measuring the Acute Toxicity of Effluents and Receiving Waters to Freshwater and Marine Organisms. Office of Water, U.S. Environmental Protection Agency, Washington, D.C., p. 33.

51. Üveges, B., Mahr, K., Szederkényi, M., Bókony, V., Hoi, H., Hettyey, A., 2016. Experimental evidence for beneficial effects of projected climate change on hibernating amphibians. Scientific Reports 6, 1–7. 10.1038/srep26754

52. van den Berg, H., da Silva Bezerra, H.S., Al-Eryani, S., Chanda, E., Nagpal, B.N., Knox, T.B., Velayudhan, R., Yadav, R.S., 2021. Recent trends in global insecticide use for disease vector control and potential implications for resistance management. Scientific Reports 11, 1–12. 10.1038/s41598-021-03367-9

53. van den Berg, H., Zaim, M., Yadav, R.S., Soares, A., Ameneshewa, B., Mnzava, A., Hii, J., Dash, A.P., Ejov, M., 2012. Global trends in the use of insecticides to control vector-borne diseases. Environmental Health Perspectives 120, 577–582. 10.1289/ehp.1104340

54. van Gelder, J.J., Olders, J.H.J., Bosch, J.W.G., Starmans, P.W., 1986. Behaviour and body temperature of hibernating common toads *Bufo bufo*. Ecography 9, 225–228. 10.1111/j.1600-0587.1986.tb01212.x

55. Vanzetto, G. V., Slaviero, J.G., Sturza, P.F., Rutkoski, C.F., Macagnan, N., Kolcenti, C., Hartmann, P.A., Ferreira, C.M., Hartmann, M.T., 2019a. Toxic effects of pyrethroids in tadpoles of Physalaemus gracilis (Anura: Leptodactylidae). Ecotoxicology 28, 1105– 1114. 10.1007/s10646-019-02115-0

56. Vanzetto, G. V., Slaviero, J.G., Sturza, P.F., Rutkoski, C.F., Macagnan, N., Kolcenti, C., Hartmann, P.A., Ferreira, C.M., Hartmann, M.T., 2019b. Toxic effects of pyrethroids in tadpoles of *Physalaemus gracilis* (Anura: Leptodactylidae). Ecotoxicology 28, 1105– 1114. 10.1007/s10646-019-02115-0

57. While, G.M., Uller, T., 2014. Quo vadis amphibia? Global warming and breeding phenology in frogs, toads and salamanders. Ecography 37, 921–929. 10.1111/ecog.00521

58. Young, H.S., McCauley, D.J., Galetti, M., Dirzo, R., 2016. Patterns, Causes, and Consequences of Anthropocene Defaunation. Annual Review of Ecology, Evolution, and Systematics 47, 333–358. 10.1146/annurev-ecolsys-112414-054142

59. Zhou, J., Zhao, H., Chen, L., Xing, X., Lv, T., Yang, X., Wu, Q., Duan, J., Ma, H., 2019. Effect of exposure to deltamethrin on the bufadienolide profiles in *Bufo bufo gargarizans* venom determined by ultra-performance liquid chromatography-triple quadrupole mass spectrometry. RSC Advances 9, 1208–1213. 10.1039/C8RA07871H

