## Supplementary material for "Effects of mild winter conditions and two pyrethroid insecticides on the development and survival of juvenile common toads (*Bufo bufo*)": Table S1

**TABLE S1:** Number of juvenile common toads introduced into outdoor enclosures, and the number of recaptured individuals. Each enclosure contained members of a given treatment group, but individuals from different families were mixed equally prior to exposure. The fence of enclosure five was damaged, so that predators could enter and the toadlets could escape.

| Treatment | Enclosure ID | Number of introduced animals | Number of recaptured animals | Number of overwintering animals |
|---|---|---|---|---|
| Control | 1 | 56 | 54 | 18 |
| | 6 | 56 | 33 | 18 |
| | 8 | 56 | 36 | 18 |
| Cypermethrin | 3 | 56 | 37 | 27 |
| | 5 | 56 | 0 | 0 |
| | 7 | 56 | 36 | 27 |
| Deltamethrin | 2 | 56 | 18 | 18 |
| | 4 | 56 | 31 | 18 |
| | 9 | 56 | 29 | 18 |
| | | | **Total: 274** | **Total: 162** |

**FIGUER S1**: Quantities of fat reserves as categorized using light microscopy. Barplots represent the number of individuals within the given category.

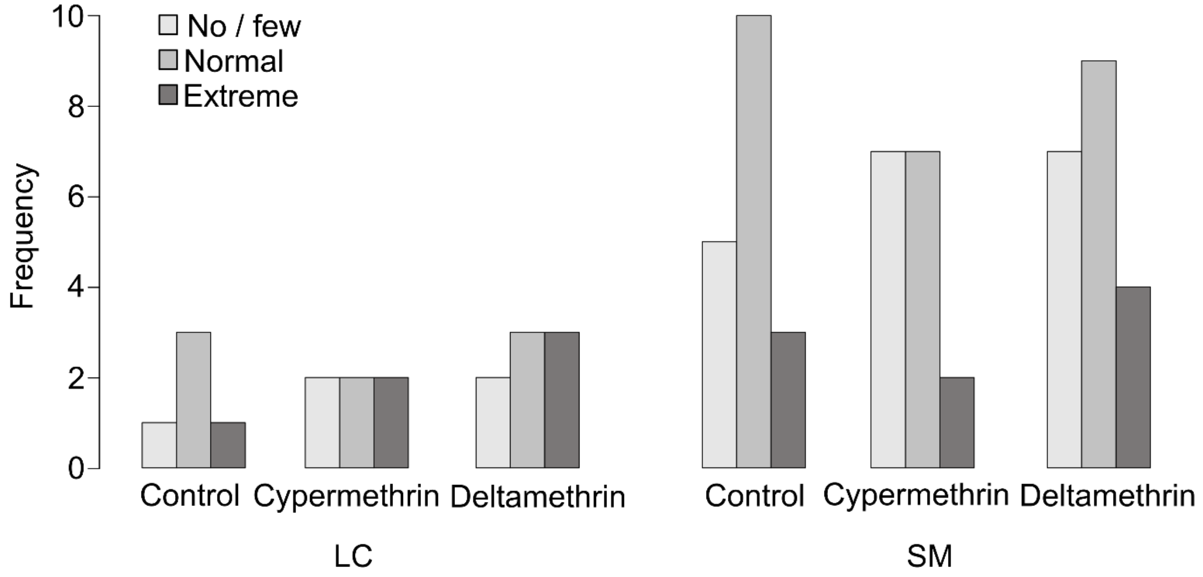
